# Rational Design of Monomeric IL37 Variants Guided by Stability and Dynamical Analyses of IL37 Dimers

**DOI:** 10.1101/2024.02.06.579100

**Authors:** Inci Sardag, Zeynep Sevval Duvenci, Serkan Belkaya, Emel Timucin

## Abstract

IL37 plays important roles in the regulation of innate immunity and its oligomeric status is critical to these roles. In its monomeric state, IL37 can effectively inhibit the inflammatory response triggered by IL18 through binding to the IL18 receptor *α*, a capability lost in its dimeric form. This paradigm underscores the pivotal role of IL37’s dimer structure in the design of novel anti-inflammatory therapeutics. Hitherto, two IL37 dimer structures were deposited in PDB, reflecting the potential use of their binding interface in the design of IL37 variants with altered dimerization tendencies. Inspection of these static structures suggested a substantial difference in their dimer interfaces. Prompted by this discrepancy, we analyzed the PDB structures of IL37 dimer (PDB: 6ncu and 5hn1) along with a predicted structure by AF2-multimer by molecular dynamics (MD) simulations to unravel whether and how IL37 can form homodimers through distinct interfaces. Results showed that the 5hn1 and AF2 dimers, which shared the same interface, stably maintained their initial conformations throughout the simulations whilst the recent IL37 dimer (PDB ID: 6ncu) with a different interface, did not. These findings underscored that the recent IL37 dimer (6ncu) structure is likely to contain an error, probably in its biological assembly record, otherwise it was not a stable assembly *in silico*. Next, focusing on the stable dimer structure of 5hn1, we have identified five critical positions of V71/Y85/I86/E89/S114 that would altogether reduce dimer stability without affecting the monomer fold. Two quintet mutations were tested similarly by MD simulations and both mutations showed either partial or complete dissociation of the dimeric form. Overall, this work contributes to the development of IL37-based therapeutics by accurately representing the dimer interface in the PDB structures and identifying five potential substitutions to effectively inhibit the inflammatory response triggered by IL18.

## 1. Introduction

Members of the interleukin-1 (IL1) family are involved in the regulation of innate immunity and inflammation (1). While the cytokine family is primarily linked to proinflammatory functions, certain members have also been identified to play anti-inflammatory roles (2; 3). IL37 stands out among other anti-inflammatory members of IL1 family because it acts in distinct pathways to reduce inflammation. Specifically, it can block inflammation intracellularly through SMAD3 interaction whilst it can also block the pro-inflammatory response led by IL18 through binding to its receptor *α* (IL18R*α*) extracellularly (4; 5). Given these critical functions, IL37 has been recognized as fundamental player of the regulation of innate immunity (6).

IL37 forms a homodimer structure (6) that negatively regulates its anti-inflammatory role by blocking its binding to the IL18R*α* (7). Thus, the monomeric form of IL37 exhibited greater efficacy in inhibiting inflammation compared to the dimeric form (6). This perspective consolidates the importance of dimer structure of IL37 and its dimer interface for desing of IL37-based therapeutics. There are two PDB structures found for IL37 both of which are annotated as homodimer. The first structure (PDB ID: 5hn1) covers the residues from 53 to 206 (6) while the recent structure (PDB ID: 6ncu) covers slightly longer region encompassing the residues 46 to 206 (8). IL37 in both of the PDB structures has *β*-trefoil fold with 12 *β* strands and 3 *α* helices (6). Despite high similarity in the 3D fold of IL37 monomers, the dimer conformations in these static structures differ significantly. Considering the significance of the IL37 dimer structure in designing potent anti-inflammatory therapies (6; 8), addressing whether or not this discrepancy holds biological significance is crucial.

Here, we have extensively investigated the IL37 dimer structures by molecular dynamics (MD) simulations and pursued a design of IL37 that would not dimerize and maintain a stable monomeric state. We have conveyed that the apparent destabilization of the recent PDB structure in the simulations may be attributed to a potential misannotation issue in its biological assembly record. Furthermore, we undertook the design of two monomer-locked variants and successfully identified a promising variant featuring five substitutions. Overall, resolving the discrepancy in the PDB dimer structures of IL37 enhances our understanding of the molecular mechanism driving the anti-inflammatory action of IL37, thereby advancing the potential for developing IL37-based therapeutics.

## 2. Methods

### 2.1. Dimer Structures

IL37 homodimer structures with the PDB IDs 6ncu and 5hn1 were extracted from PDB. Modeling of the missing regions for both dimer structures were performed by MODELLER plugin (9) of ChimeraX (10). Several loop models were generated and the final models were selected according to the lowest normalized Discrete Optimized Protein Energy (zDOPE) score, for which a negative score signifies better predictions (11). Homodimer structure of IL37 was also predicted by AlphaFold2 multimer (AF2) (12; 13; 14; 15). MSAs were performed by MMseqs2 software (16; 17) with no template mode. MSA mode was selected as MMseqsUniRef + Environmental whereas pair sequences from the same species and unpaired MSA was chosen as the pairing mode (18). The number of cycles was adjusted to 48 (19). Predicted structure was relaxed by AMBER force field to resolve clashes (20). The final model was selected based on residue confidence score (pLDDT), which relies on the lDDT-C*α* metric (21), pairwise error prediction score (pTM) and predicted aligned error (PAE).

### 2.2. Molecular Dynamics Simulations

Three IL37 dimer structures composed of two PDB (6ncu and 5hn1) and one AF2 computed structure were analyzed by molecular dynamics (MD) simulations. All MD systems were prepared using CHARMM-GUI (22; 23; 24). Crystal water and other solvent molecules were removed and the structures were protonated according to pH 7.0. Complexes were placed in the center of rectangular boxes with 10 Å of edge distances, which were fit according to size of the complexes. Na^+^ and Cl*^−^* counterions were placed to neutralize the systems to a final concentration of 0.15 M. MD simulations were carried via the NAMD engine (25) using the CHARMM36m force field with a *ϕ*, *ψ* grid-based energy correction map (26; 23; 27). Water molecules were treated explicitly by the TIP3P model (28). Periodic boundary conditions and a time-step of 2 fs were applied to all simulations. Particle-mesh Ewald method was applied for computation of long-range electrostatic interactions with a grid spacing of 1 Å (29). A cut-off distance of 12 Å was used for the non-bonded interaction terms. Three IL37 homodimer systems were energy-minimized in 10,000 steps and equilibrated for 250 ps in an *nVT* ensemble. Finally, systems were simulated for 500 ns under constant pressure (1 atm) and temperature (310.15 K) using the *nPT* ensemble using the Langevin thermostat and piston pressure method (30; 31; 32). Production simulations were repeated three times for each system.

### 2.3. Trajectory Analysis

Production MD trajectories were analyzed by means of pairwise root mean square displacement (RMSD) and fluctuation of C*α* atoms (RMSF). For the former, MDanalysis scripts were used (33; 34). Time-dependent change in the solvent accessible surface area (SASA) of the dimer and subunits was monitored. Interface SASA was calculated as the difference in SASA measurement between the sum of each subunit and dimer. Visualization of the structures and trajectories was performed by Visual Molecular Dynamics (VMD) (35). Bio3D package was used for the principal component analysis (PCA) (36). SASA and fluctuation calculations were conducted by the *measure* command of VMD (35).

### 2.4. FoldX Calculations

Two snapshots were selected from each replicate trajectory, one from the last 10 ns of the first half of the trajectory and the second from the last 10 ns of the second half. Each system was represented by a total of seven snapshots including the static structure (PDB or AF2 computed structure). Each snapshot was then analyzed by the protein design tool, FoldX (37). The FoldX command *AnalyseComplex* was initially run to gather the interface amino acids for the 5hn1 structure. This list of interface amino acids were then analyzed by the *Pssm* command to calculate their contribution to the binding free energy of the dimer. Site-saturation mutagenesis were considered for each selected position. Lastly, the command *AlaScan* was used to map the contribution of the IL37 amino acids to the folding free energy of the dimers and subunits.

## 3. Results

### 3.1. Static Structures of Human IL37 Dimer

IL37 dimer structure was captured in two crystal structures (PDB IDs: 6ncu and 5hn1) (8; 6). Fig. 1a-b depicts the biological assemblies of these structures. A cyclic C2 symmetry was described for the biological assembly of the 5hn1 (6), which refers to a rotation of 180*^◦^* around a line. However, no recognizable symmetry elements were found for the 6ncu structure (8). Close-up views of the dimer interfaces also reveal the conflict in the dimer configurations of two assemblies. Specifically, the 5hn1 dimer interface was formed by a symmetrical contour while it was not the case for 6ncu dimer. Two symmetrical electrostatic interactions between K83-D73 link two subunits at the poles of the binding interface of 5hn1 (Fig. 1a), leaving a hydrophobic cluster in the interior. However, 6ncu, both as a biological assembly and asymmetric unit, neither shared this interface nor had a partial overlap in its interface with the 5hn1 dimer (Fig. 1b). Instead, the dimer interface of 6ncu contoured the asymmetric surfaces of the IL37 monomers, resulting in a less compact interface without any detectable network of interactions. Superimposition of both dimers confirmed this difference in the biological assemblies of these two structures (Fig. 1c). The superimposition of both dimers relying on only chain A and chain B, reflecting that 6ncu dimer interface is asymmetrically formed by distinct regions of the subunits (Fig. S1). Furthermore, the reported interface in the 6ncu dimer was quite different than that of 5hn1 dimer (Fig. S2). Overall, this comparative analysis of two biological assemblies from PDB implies two different dimer binding modes for the IL37 monomers.

**Figure 1:**
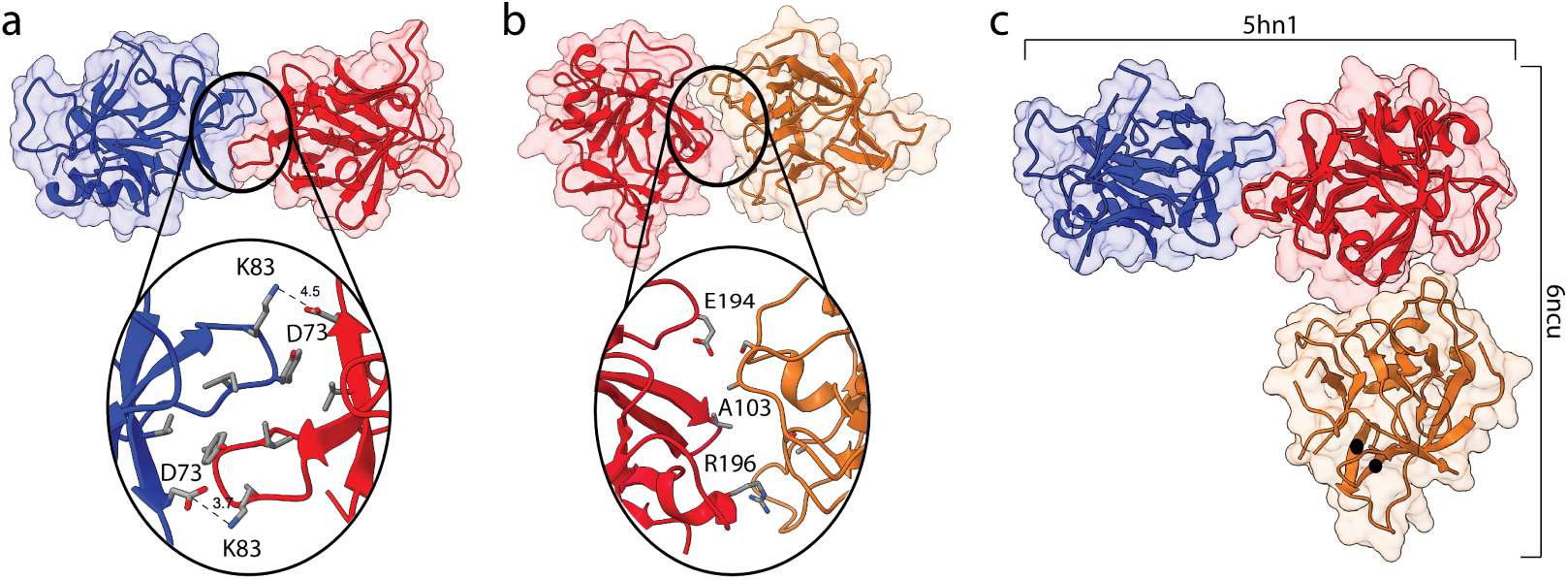
Binding interface of two crystal dimers 5hn1 and 6ncu from PDB. (a) Hydrophobic interface of the 5hn1 dimer composed of V71, Y85 and I86 from both subunits, which is sectioned by two symmetrical intermolecular interactions between K83 and D73, is visualized. (b) The interaction surface of 6ncu is depicted showing the amino acids that are closely found at the interface. (c) Best superimposition of two biological assemblies were shown based on C*α* RMSD.

The IL37 homodimer was also predicted by AF2 multimer (Fig. S3), showing a highly accurate prediction based on pLDDT and predicted aligned error (PAE) scores. Superimposition of the 5hn1 and AF2 dimer structures in (Fig. S3) showed that AF2 model adapted the same dimer configuration with the 5hn1 structure with an average change in C*α* RMSD of 1.3 Å. The overall structures and binding interfaces of the 5hn1 and AF2 predicted dimers were in close agreement, except for two disordered regions exhibiting notably low pLDDT scores, which contributed to an increase in the RMSD. We also calculated the surface area of the dimer interfaces in these structures (Table S1). Notably, 6ncu dimer interface was much smaller than the 5hn1 and AF2 predicted dimers. Overall, detailed analysis of these static dimer structures revealed an apparent discrepancy in the PDB structures of 6ncu and 5hn1 as well as in the AF2 computed dimer. Considering the crucial role of the IL37 dimer structure in the design of effective anti-inflammatory therapies (8), addressing this inconsistency is imperative.

### 3.2. Dimer Dynamics and Stability

After modeling of the missing regions of PDB structures, we have analyzed all three IL37 dimers by all-atom MD simulations. Details of the MD systems were given in Table S2. All simulations were lasted for 0.5 *µ*s and repeated three times. Pairwise C*α* RMSD was calculated for all dimers (Figure S4). The 5hn1 and AF2 predicted dimers showed no significant C*α* mobility when the dimer backbone was set as the reference, while the 6ncu dimer backbone has reached RMSD values as high as 20 Å for all three simulations (Figure S4a). To identify whether this large mobility of 6ncu is due to monomer or dimer instability, we extended the RMSD calculations using only single monomer structure as the reference. For all three dimers, the chain A stayed intact throughout all simulations (Fig.S4b-left). For the 5hn1 and AF2 dimer structures, a slight increase was reported in the C*α* mobility of chain B with respect to chain A, the observation which was expected given the reference alignment on the chain A (Fig.S4b-right). However, the 6ncu dimer showed a highly mobile chain B with respect to the chain A for all simulations (Fig.S4b), suggesting disruption of the dimer structure.

We also extracted essential dynamics of each system by cartesian PCA (Fig. S5). Score plots obtained from the first three principal components (pcs) corroborated that 6ncu dimer spanned a much larger conformational space than the other dimers (Fig. S5). Altogether, these results suggested that highly mobile 6ncu dimer was likely a result of dimer instability rather than monomer instability, whereas for the 5hn1 and AF2 structures, neither dimer nor monomers displayed an abnormal level of backbone mobility that could be an indication of instability.

We also examined the solvent accessible surface area (SASA) of the dimer, monomers, and interface for all three systems (Fig.2). The SASA profiles of the dimer and monomers displayed stabilization in the case of the 5hn1 and AF2 dimers. However, in the 6ncu structure, particularly the dimer and interface SASA exhibited significant fluctuations. Notably, the interface SASA was measured to be zero for one of the simulations of 6ncu (Fig.2, lightest green), indicating complete dissociation of the dimer. Nevertheless, the other dimers sharing the same interface demonstrated a stable interface, as evidenced by flat SASA measurements of the interface (Fig. 2).

**Figure 2:**
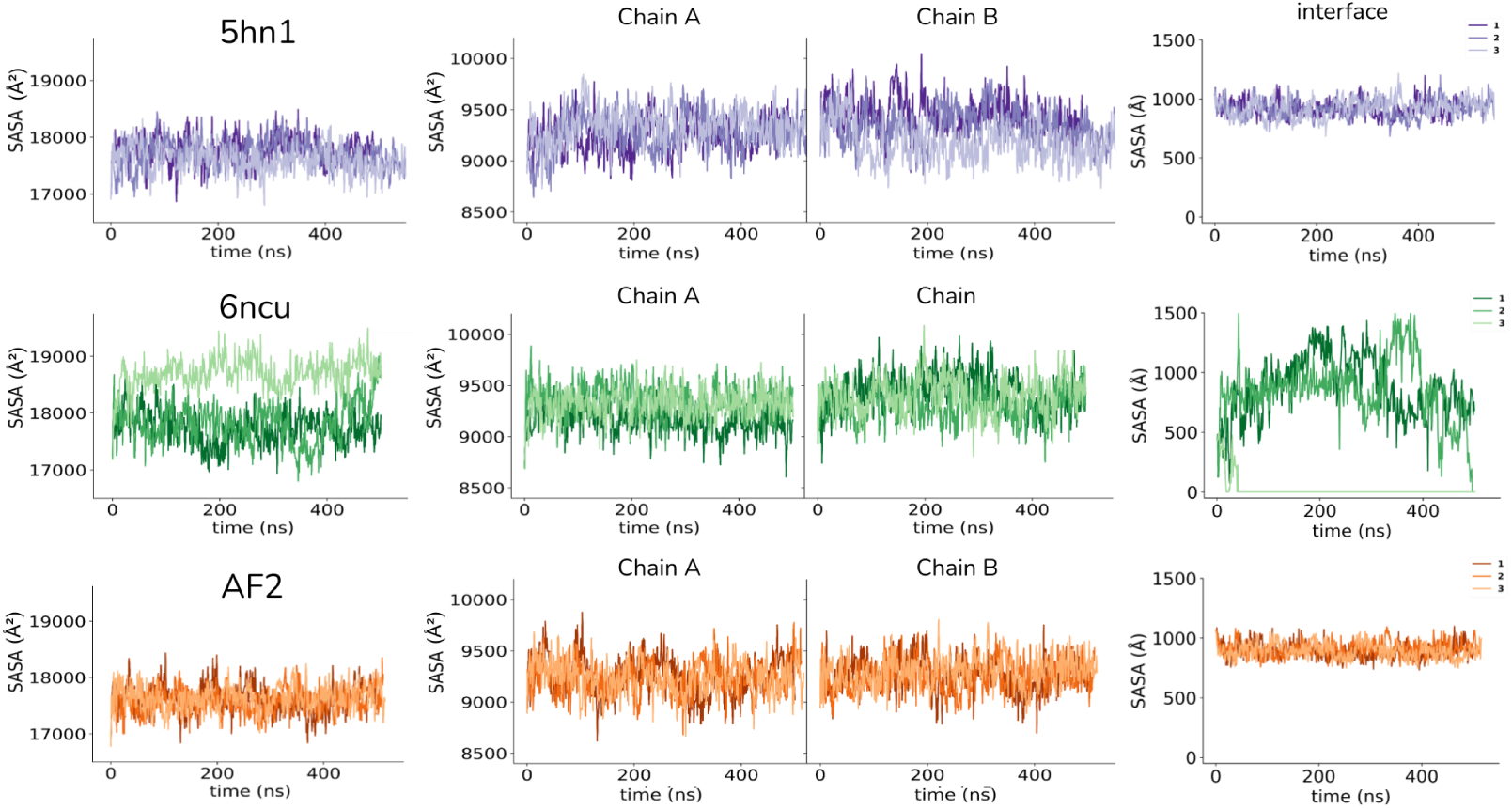
Solvent accessible surface area (SASA) changes were calculated for all three dimers. First column shows the dimer SASA, the second and third columns show monomer SASA values, and the last column shows the interface SASA calculated by subtracting the sum of subunit SASA values from the dimer SASA.

To further investigate how the overall dimer structure has changed, reduced trajectory of the first simulations was visualized for each system (Fig. 3). These reduced trajectories show the initial and final structures of the chain B in red and blue colors respectively and also shows the shortened trajectory of the center of mass of each subunit using a sphere representation. Given this analysis, there is almost no relative change in the orientation of subunits in the 5hn1 and AF2 dimers. However, the 6ncu dimer showed a dramatic movement of chain B with respect to chain A, indicating the disruption of the dimer conformation reported in the crystal structure.

**Figure 3:**
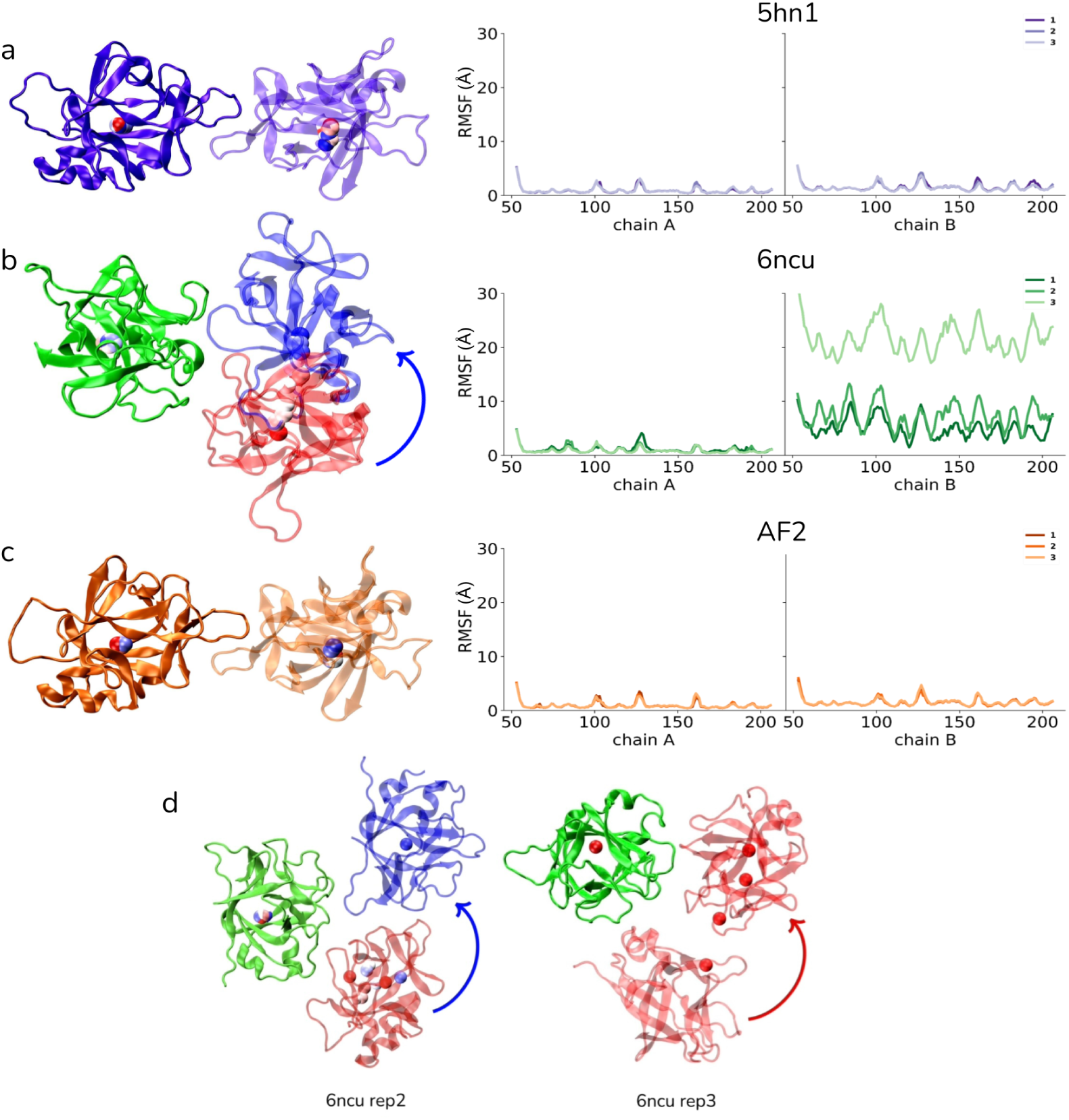
The relative displacement of one monomer (chain B) with respect to the other (chain A) for the first simulations of the (a) 5hn1, (b) 6ncu and (c) AF2 dimers. For each visual, center of mass of the monomers were shown by spheres colored according to simulation time (red: start, blue: end). (d) Panel shows the rest of replicates simulations of 6ncu, wherein the dimer subunits were almost completely dissociated.

Right panel in Fig. 3 demonstrates C*α* fluctuations for each monomer, wherein no major fluctuations were observed for the 5hn1 and AF2 dimers. However, all residues in one of the monomers of the 6ncu dimer showed much higher fluctuations than those in the other monomer, indicating that one of the monomer in 6ncu has been displaced with respect to the other chain (Fig. 3b-d).

We also traced SASA of each subunit interface in all systems (Fig. 4). First, we noted that the PDB conformations of the subunit interfaces formed by the continuous epitope between residues 80-87 perfectly aligned with each other in two crystal structures, 5hn1 and 6ncu. However, during the simulations, this surface epitope in the 6ncu subunits underwent notable changes that resulted in an extension of this surface, disrupting the hydrophobic cluster mediated by Y85. On the other hand, two other structures, 5hn1 and AF2-computed dimers did not show any apparent change in shape or SASA of this continuous epitope forming the dimer interface. This analysis particularly pointed out that the subunit interface between 80-87 was highly flexible in the 6ncu dimer, undergoing conformational changes that would limit this epitope’s interaction with the other subunit.

**Figure 4:**
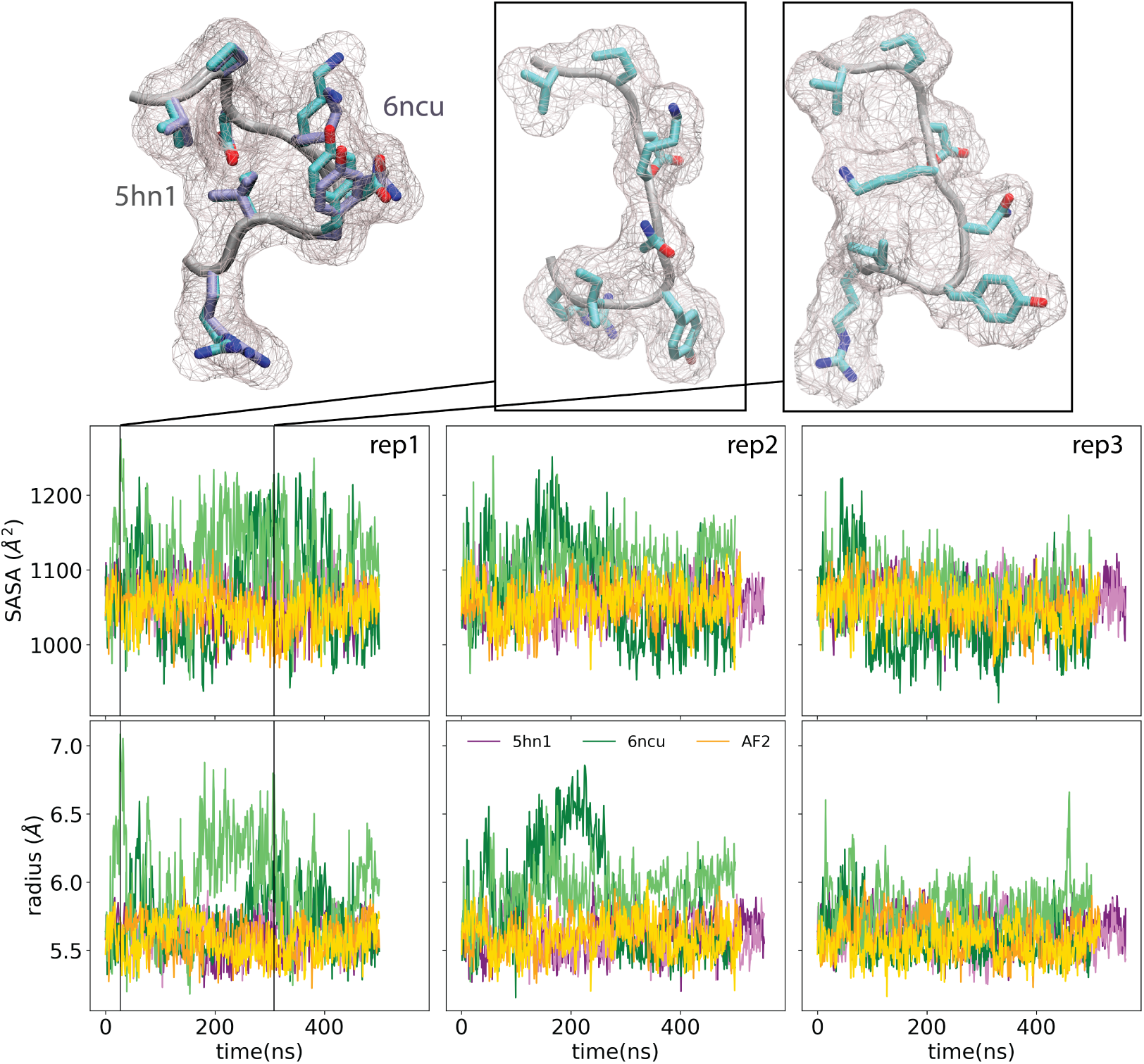
Superimposed dimer interface (80-87) of 5hn1 (cyan) and 6ncu (grey) structures were shown. SASA and radius of gyration (*R_G_*) of this interface was measured throughout simulations for both chains. 6ncu chains were colored by green, 5hn1 chains were colored by orange and AF2 chains were colored by purple. Top row contains two representative snapshots from the 6ncu simulations showing a notable change in the SASA and *R_G_* of the epitope.

We lastly conducted an *in silico* alanine scanning experiment by FoldX for all three IL37 dimer structures. A selected set of seven snapshots including the PDB structure and two snapshots from each repeated simulations were recruited to calculate the average change in stability (ΔΔ*G*) upon alanine mutation of each position in the IL37 dimer. Essentially, alanine substitution at the dimer interface positions of the 6ncu dimer resulted in negligible changes in the stability. On the other hand, alanine substitutions of the dimer interface of the 5hn1 and AF2 structures destabilized the structure by over 2 kcal.mol^-1^. This analysis further revealed the presence of additional interactions in the 5hn1 and AF2 dimers, which were absent in the 6ncu dimer. Overall, the extensive analysis of all-atom MD simulations underscored that among two distinct dimer conformations reported in the PDB, only 5hn1 is a stable dimer.

### 3.3. Human IL37 Variants and Their Impact on Stability

We have next extracted all reported human missense variations of the mature IL37 from gnomAD v3.1.2 (38) with the ID of ENST00000263326.8. The structural impact of these variants on the monomer and dimer stability was investigated by FoldX (37) (Table S3). Only two of these variants, I177T and R152W were annotated with a clinical significance label as pathogenic and benign, respectively. Essentially, these two positions did not make any close contacts with the other subunit, i.e. they are not found at the dimer interface. In line with this, the substitutions of I177T and R152W do not affect dimer stability at all, as reflected by the ΔΔ*G_dimer_* scores (Fig. 5a). However, both substitutions caused a destabilization impact on monomer, with a positive ΔΔ*G_monomer_* score greater than 2 kcal.mol*^−^*1. Although I177T and R152W had a similar magnitude of a destabilization effect, I177T was pathogenic and associated with inflammatory bowel disease (39) whilst R152W was a benign variant according to ClinVar. This discrepancy could be have been due to the relative accessibility of the positions in question. Particularly, I177T was located at the core and its side chain was not accessible. On the other hand, R152 was not completely buried and much more accessible than I177 (Fig. 5a). Hence, substitution of R152 to another bulky amino acid of W might impact the core structure of IL37 less than did I177T substitution.

**Figure 5:**
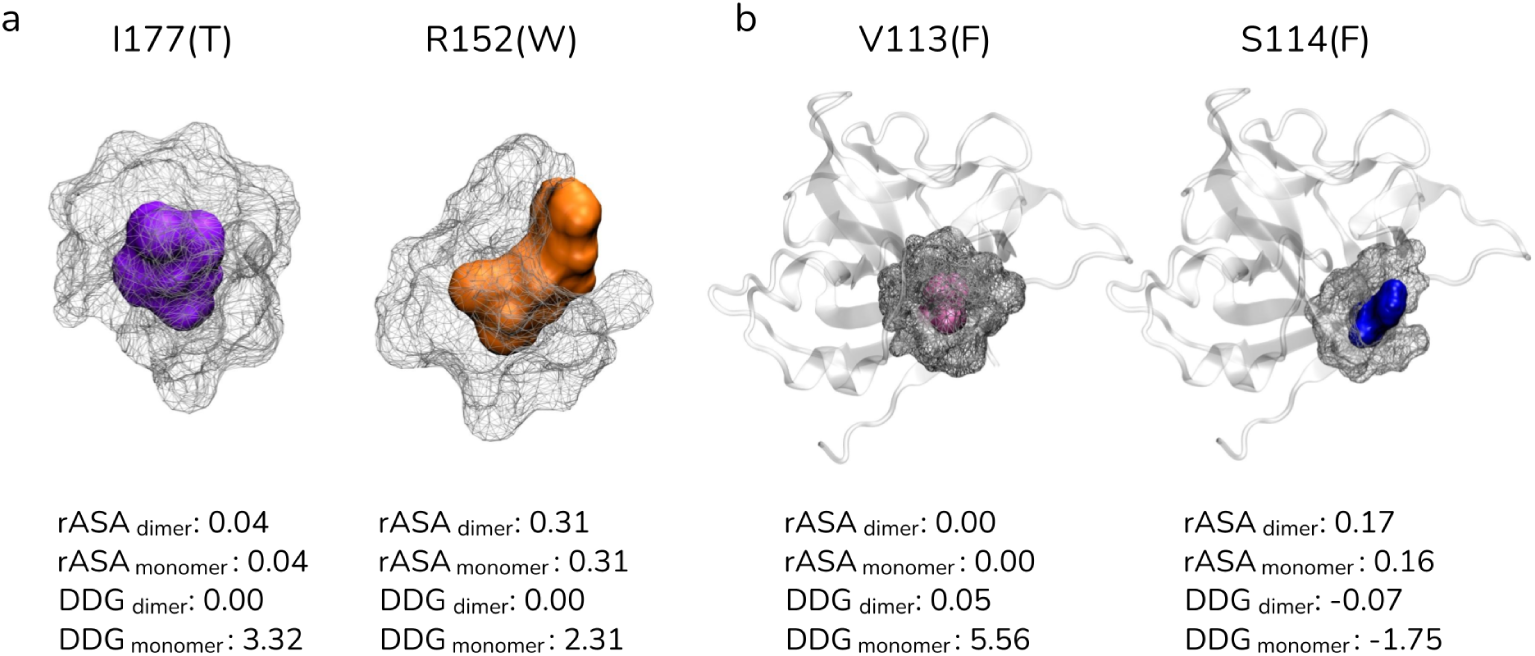
The locations of selected human IL37 variants, I177T, R152T, V113F and S114F and other residues within range 5 Å. (a) The positions I177R152 and (b) and (b) V113 and S114 along with surrounding amino acids were shown in surface representations. Relative accessible surface area for the dimer and monomer structures were also given for all amino acids. The impact of the observed human variants of these positions on the dimer and monomer stability were given in kcal.mol^-1^.

We also point out a similar case for two other substitutions V113F and S114F whose FoldX calculations led to contradictory scores though they are adjacent positions and substituted to same amino acid (Fig. 5b). Particularly, V113F led to a notable decrease in IL37 stability by 5.6 kcal.mol^-1^, while S114 led to a slight increase in the stability by -1.8 kcal.mol^-1^ (Fig. 5b). This large difference could be linked to the relatively higher accessibility of S114 compared to the V113. The V133F, albeit being a complementing substitution from V to F, would disrupt the tight interactions formed by V113. Thus, V113F might be clinically more important given its parallel outcome with I177 in FoldX analysis (Fig. 5a).

### 3.4. Design of Novel IL37 Variant with Reduced Dimer Stability

Novel and reported variants of IL37 homodimer that might destabilize dimer structure without interfering with the monomer structure were evaluated. Seven snapshots of 5hn1 and AF2 that were obtained from repeated MD simulations were employed in this analysis. We carried out *in silico* site-saturation mutagenesis for each selected position and analyzed their impact on the stability of dimer (Fig. S7) and monomer forms (Fig. S8).

We pursued substitutions that yielded positive scores in the binding free energy of the dimer (ΔΔ*G_dimer_*), which is an indication of dimer destabilization (Fig. S7) and negative scores in the folding free energy of the monomer (ΔΔ*G_monomer_*), as an indication of monomer stabilization (Fig. S8). Any substitution at the positions of Y85 and I86 showed destabilization of dimer by more than 1 kcal.mol^-1^ (Fig. S7), confirming their pivotal involvement in the dimerization (40). Together with these extensive analyses by FoldX, we have reported three additional positions of V71, E89 and S114 that would impair dimer structure without significantly affecting the monomer stability. Overall, the mutations of V71R, Y85K/C, I86W were selected as the most promising dimer stabilizing mutations (Fig. S7) with small destabilization of the monomer (Fig. S8). Furthermore, E89L and S114R mutations were chosen because of their stabilizing impact on monomer structure (Fig. S8). Overall, two quintet-mutants were generated including the V71R/Y85C/I86W/E89L/S114R and V71R/Y85K/I86W/E89L/S114R mutations and varying only for the Y85 substitution. We pursued both a cysteine and an arginine substitution at the Y85 position as extensive FoldX analysis underscored a large dimer stabilizing impact for both substitutions (Fig. S7). Particularly, Y85C destabilized the dimer structure by 2.91 kcal.mol^-1^ and the monomer by 0.92 kcal.mol^-1^. Similarly, Y85K destabilized the dimer structure by 2.38 kcal.mol^-1^ and the monomers by 0.54 kcal.mol^-1^. Both of the quintet mutants have overall destabilized dimer by 6.52 and 8.430 kcal.mol^-1^. This pronounced dimer destabilization impact in the quintet mutations showed the destabilization effect of the single mutations were added-up rather than cancelled out.

Both quintet mutants were analyzed in MD simulations similarly in the dimeric form. Both simulations showed an immediate disruption of the dimer interface, reflecting a significant change in the resulting conformations with respect to the wild-type dimer (Fig. 5). Specifically, the Y85C containing mutant caused a complete dissociation of the dimer in 29.0 ns while the mutant containing Y85K led to a partial dissociation of the dimer in 50.8 ns (see supplementary movies). We observed an interaction between Y85K of one monomer and D73 of the other monomer, in part explaining the partial dissociation of this mutant. Besides its dimer destabilization impact, the Y85K substitution also destabilized the monomer (Fig. 5). On the other hand, the mutant containing Y85C exclusively destabilized the dimer structure without significantly affecting the monomer stability. Overall, the quintet mutation harboring Y85C has the most optimal ratio of dimer-to-monomer stability, while this ratio was not desirable for the quintet mutant having Y85K substitution (Fig. 5). Hence, the quintet mutation holding Y85C is proposed to be a favorable variant to effectively disrupt the dimer structure without affecting the monomer stability.

## 4. Discussion

Dimerization of proteins, either in hetero- or homo-dimeric form, is an essential regulatory process that affects their biological function (41; 42). Head-to-head conformation is one of the homodimer arrangements addressing to the relative positions of the subunits based on the evolution of symmetry (43). The N terminus of one monomer is in close proximity with the N terminus of another monomer in head-to-head conformation (43). Head-to-head dimerization is a common among IL1 protein family cytokines (6). Although dimerization mostly blocks the biological activity of IL1 members, i.e. pro-inflammatory response, the opposite is the case for IL37 as it can achieve its anti-inflammatory role in the monomeric state (6). Hence, IL37 engineering approaches creating stable monomeric variants of IL37 is recognized to be a potential avenue for anti-inflammatory therapies (6). This recognition underscores the importance of the dimer structure of IL37 that would template the design process.

The deposited IL37 structures in PDB were both annotated as homodimers, albeit they have different dimer conformations (Fig. 1). Alongside a notable distinction in their dimer interfaces, we also noted that the dynamics of these dimers differ a lot (see supplementary movies). Particularly, the recent structure (6ncu), which did not have any symmetry elements but formed through an asymmetrically countered dimer interface, was not stable and consistently dissociated, either partially or completely. On the contrary, the first PDB structure (5hn1) and the AF2-computed dimer with a highly similar dimer interface stayed intact without any peaked mobility or flexibility that could be suggestive of dimer instability.

An additional noteworthy observation regarding the static dimer interfaces is the complete inaccessibility of residues at the 5hn1 dimer interface to the solvent. These residues form airtight interactions with the other subunit, as illustrated in Figure 1c. Conversely, the interface of 6ncu is porous, which would allow water molecules to penetrate (Fig. 1c). This disparity in the compactness of the dimer interfaces contributes to a stability difference in the dimers (44), thereby in part explaining the lower stability of 6ncu compared to 5hn1.

The discrepancies spotted in the static and dynamic structures of the dimers may stem from various factors. It is plausible that misannotation issues in the 6ncu structure or errors in the deposition of its coordinates are responsible for these differences. Otherwise, our simulations did not include any user-defined forces or the dimer structures were not extensively manipulated, that could result in the suboptimal dynamics for the 6ncu dimer in our observations. The only manipulation of the PDB structures was the modeling of missing regions, corresponding to two very short stretches (3-, 1-amino acid long) for the 6ncu structure (8). While the missing regions in the 5hn1 structure that were relatively longer than those in the 6ncu, were also modeled, underlining a similar treatment for both PDB dimers before simulations.

Notably, the crystal structure of 5hn1 was captured in the presence of 158 water and 5 sulfate molecules (6), whereas there were not any crystal waters in the 6ncu structure. Thus, stripping of crystal waters, which might have an effect on the simulation outcome (45; 46), was not experienced by the 6ncu structure. Water is essential for proteins to maintain their native structural conformations during crystallization (47). Tightly bound immobile water molecules that form the first hydration shell around the protein are expected to be present in many PDB entries (48). The proportion of crystal protein volume occupied by the aqueous solvent ranges from 27% to 65%, with an average fraction of 43% (49). Therefore, PDB entries with less crystal waters than 5% or more than 90% tend to indicate serious errors about the protein structure and wrong annotations (48). Hence, we reported that the absence of crystal waters in the 6ncu structure may also indicate an issue with this deposition.

Erroneous annotation of the biological unit of PDB structures is an important source of error relating to the experimental solution state of the protein structure (50; 51; 52). The error rate in biological unit annotations were estimated in the range of 7-15% (53; 51). These errors can sourced from various factors such as biases during model building, poor resolution, crystallization conditions and/or biological complexity (51; 50). Furthermore, Xu et al. have highlighted some cases wherein the biological assembly annotation of the PDB structure, which was provided by the depositors in the REMARK 350, differs from what has been described in the linked publication (50; 51). In this sense, the 6ncu is likely to be an example to such cases, because the IL37 homodimer structure of 6ncu was described as being almost identical to the 5hn1 structure in its corresponding publication (8). Overall, our dimer simulations reported a comparative perspective on the disparity in the PDB structures of IL37 validated the 5hn1 structure as the stable and correct dimer structure and noted particular inconsistencies for the dimer structure of 6ncu.

Two prominent mutations of Y85A and D73K/A were found effective to block inflammation more effective than the wild-type IL37 (54). These mutations were previously introduced to disrupt the dimeric form (8; 6). However, monomer-locked IL37 structures harboring these substitutions were not reported to be effective due to rapid renal clearance due to small protein size of monomers (55; 54). Hence, IL37 engineering approaches that focus on creating variants not only with reduced dimerization potential but with an enhanced monomer stability promises to be a potential strategy for designing IL37-based anti-inflammatory therapeutics (6). Our study has utilized this strategy, computing the impact of the design on the stability of both monomer and dimer structures. Furthermore, alanine is arguably perceived as the best replacement amino acid to test the impact of a mutation (56; 57; 58); however, systematic investigation of large-scale mutagenesis libraries established that particularly charged or polar amino acids are better at disruption intermolecular interactions (59). Thus, we have applied both alanine scanning and site-saturation mutagenesis to identify dimer destabilizing and monomer stabilizing substitutions (Fig. S7-8).

Although D73 was reported to be an effective amino acid, lying at the dimer interface and forming a salt-bridge interaction with the K83 in the other subunit, our stability analysis documented that none of the D73 and K83 substitutions were critical for dimer stability (Fig. S7). Instead of the charged interactions, our analysis have underscored the significance of the hydrophobic cluster at the dimer interface for the integrity of the dimer and also of the monomer. Particularly, the stability analysis of the monomer showed that the disruption of this cluster, composed of V71, Y85, I86 and K83 -particularly the hydrocarbon sidechain of K83- (Fig. 1)a), both impairs the dimer interface (Fig. S7) and destabilizes the intramolecular interactions of the monomer (Fig. S8). Hence, any substitution at these positions should be cautiously made, even though they are intuitively prime candidates for the disruption of IL37 dimer because of being located at the dimer interface.

The Y85 plays a central role within this hydrophobic cluster for both dimer and monomer stability, as reflected by the monomer-locked status of Y85A REF. Our results showed that a cysteine or a lysine substitution was more effective in destabilizing the dimer at the Y85*^th^* position instead of an alanine substitution. Comparison of the Y85C and Y85K variants have further rendered Y85C, which was a reported human variant, as the best substitution because it prevents a additional interaction with the D73 in the other monomer, which was otherwise possible for the Y85K (Fig. 6).

**Figure 6:**
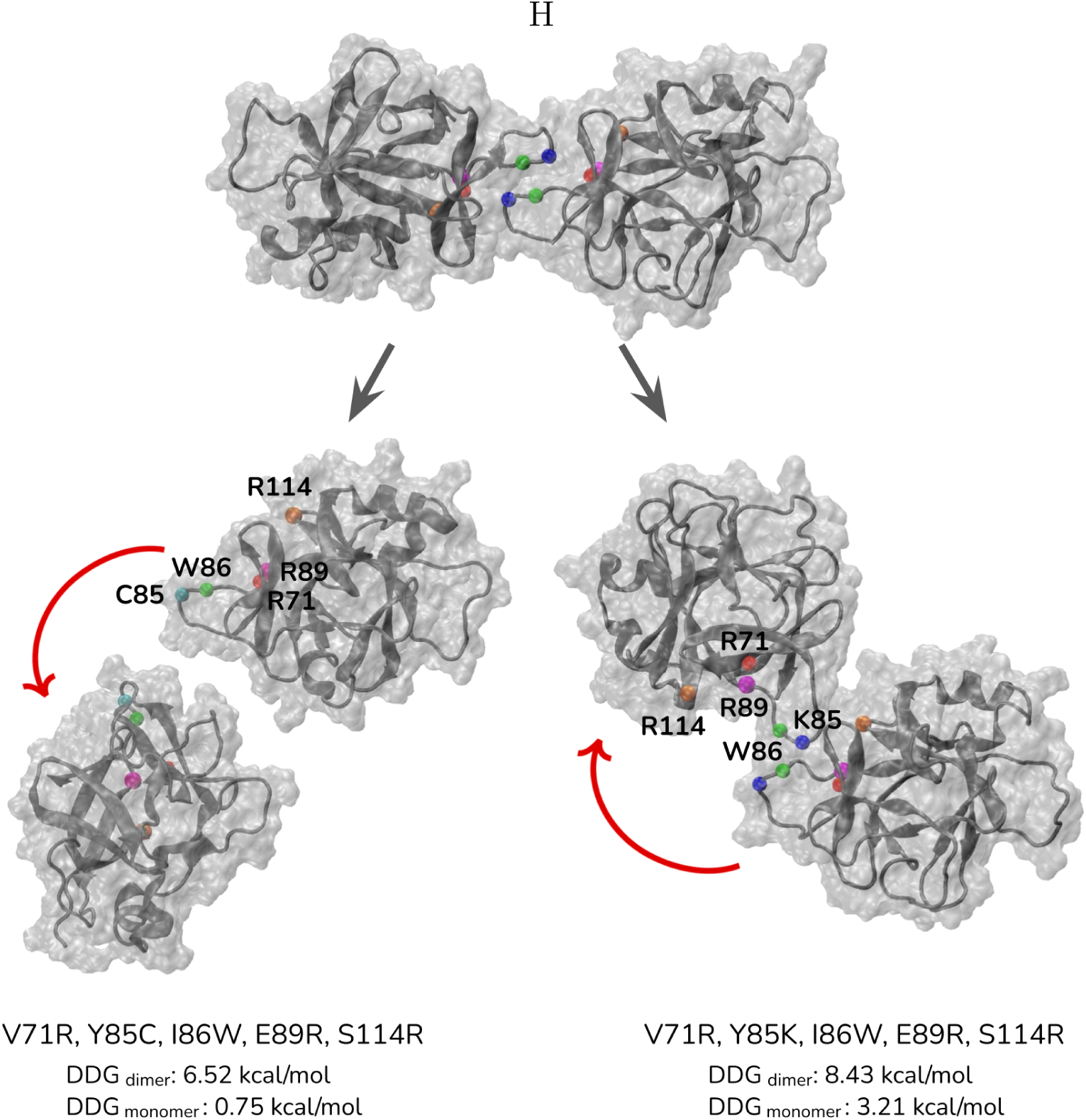
Final snapshots of the quintet mutants of IL37 after 29.0 and 50.8 ns of MD simulation of Y85C and Y85K variants, respectively. The movement of one of the monomers were shown by a red arrow, wherein the trajectory was aligned to the other monomer. The effect of these combinations on the dimer and monomer stability that were calculated by FoldX were given.

Through comprehensive analysis of seven different snapshots representing three independent MD simulations, we identified the positions of V71, I86, E89 and S114 whose substitutions would destabilize the dimer and stabilize/neutralize the monomer. Ultimately, the proposed quintet mutation of V71R/Y85C/I86W/E89L/S114R was found highly effective in disrupting the IL37 dimer without destabilizing the monomer structure *in silico* (Figs. 6, S6-8).

In conclusion, our findings (i) showed that the correct IL37 homodimer in the PDB is the 5hn1 structure while the dimer interface presented in the 6ncu structure is not stable in simulations and (ii) reported the rational design, proposing five mutations in IL37 that tune both the dimer and monomer stability.

## 5 Acknowledgements

This work was supported by the EMBO Installation Grant (#4727 to Serkan Belkaya) through the Scientific and Technological Research Council of Turkey (TÜBİTAK), Turkey.

